# Objective sleep quality predicts subjective sleep ratings: a multiday observational study

**DOI:** 10.1101/2023.12.29.573622

**Authors:** Róbert Pierson-Bartel, Péter Przemyslaw Ujma

## Abstract

In both clinical and observational studies, sleep quality is usually assessed by subjective self-report. The literature is mixed about how accurately these self-reports track objectively (e.g. via polysomnography) assessed sleep quality, with frequent reports of a very low or no association. However, previous research on this question focused on between-subject designs, which may be confounded by trait-level variables. In the current study, we used the novel Budapest Sleep, Experiences and Traits Study (BSETS) dataset to investigate if within-subject differences in subjectively reported sleep quality are related to sleep macrostructure and quantitative EEG variables assessed using a mobile EEG headband. We found clear evidence that within-subject variations in sleep onset latency, wake after sleep onset, total sleep time, and sleep efficiency affect self-reported sleep quality in the morning. These effects were replicated if detailed sleep composition metrics (percentage and latency of specific vigilance states) or two alternative measures of subjective sleep quality are used instead. We found no effect of the number of awakenings or relative EEG delta and sigma power. Between-subject effects (relationships between individual mean values of sleep metrics and subjective sleep quality) were also found, highlighting that analyses focusing only on these may be erroneous. Our findings show that while previous investigations of this issue may have been confounded by between-subject effects, objective sleep quality is indeed reflected in subjective sleep ratings.

## Introduction

Poor sleep quality is associated with several health outcomes which carry a significant burden to affected patients, health care providers, and ultimately society as a whole (Grandner, 2022). For convenience, poor sleep quality is typically assessed based on subjective self-reports (Fabbri et al., 2021). While subjectively reported sleep may provide important information about health by itself (Buysse, 2014; Utsumi et al., 2022), it is crucial to understand to what extent self-reports of sleep quality correlate with objective, instrument-based measures of sleep structure. This is because sleep complaints in the absence of objective alterations of sleep likely have different etiology and require different therapy than those which accurately reflect poor sleep (Buysse, 2014).

Several previous studies compared subjectively assessed sleep quality with objectively measured sleep macrostructure. For example, Armitage and colleagues (Armitage et al., 1997) found that in 49 healthy subjects, subjectively rated sleep quality correlated -0.5 with the polysomnography-based number of awakenings, 0.39 with slow wave sleep percentage, and 0.13 with sleep inertia. Keklund & Akerstedt (Keklund and Akerstedt, 1997) compared polysomnography-based sleep macrostructure and subjective self-reports of sleep quality in the following morning in a sample of 37 participants. They found that longer sleep, higher sleep efficiency, less time spent in wake and more time spent in slow wave sleep was significantly correlated with higher sleep quality. Recently, Gabryelska et al (Gabryelska et al., 2019) used a multivariate approach to associate power spectral density of the sleep EEG with subjective sleep quality reported the following morning. They found that weak (r∼0.1) but significant correlations exist. A very large study of over 5000 elderly American men (Unruh et al., 2008), however, found no significant correlations between sleep efficiency, arousals per hour, slow wave sleep percentage and “subjective complaints of feeling unrested, overly sleepy or not getting enough sleep”. Two more recent multivariate analyses of the same dataset including machine learning (Kaplan et al., 2017b, 2017a) found significant, but weak associations. A recent review of physiological markers of sleep quality (McCarter et al., 2022) reviewed 49 EEG-based studies to also conclude that “correlations between objectively measured sleep and objective performance or subjectively assessed sleep quality were weak to moderate”. (See also (Cudney et al., 2022) for another systematic review with a smaller number of included studies.) Thus, based on the previous literature, subjective and objective sleep quality may exhibit a small but significant correlation at best, mysteriously leaving a major part of variance in sleep quality self-reports unexplained by actual objective indicators of sleep.

These studies, however, all employed a between-subjects design. In other words, they collected a single subjective and a single objective estimate from each participant and calculated correlations between the two. This approach is problematic because these correlations may be confounded by trait-level characteristics such as personality or response tendencies. For example, it is possible that older or more depression-prone individuals systematically report worse subjective ratings of sleep even in the absence of objective alterations. This would bias the correlations of objective and subjective ratings of sleep downward, as the latter essentially reflect a mixture of actual perceived sleep quality and personality. Even in the absence of such biases, a between-subjects correlation is a poor estimate of the accuracy of self-reports, as different participants are compared instead of multiple reports from the same person reporting fluctuations of sleep quality over several nights. Ideally, self-reports of sleep quality could accurately track between-night fluctuations in objective sleep quality within the same individual.

The solution for these problems is a multiday observational study (Taji et al., 2023) (see also (Konjarski et al., 2018) for a review of similar studies). In a multiday observational study, each participant provides several assessments of both subjective and objective sleep quality and multilevel regression models are used to analyze the data. Such a design is still capable of providing between-participant estimates (essentially correlations of participant means), with the advantage that multiple assessments reduce measurement error. However, crucially, this design is capable to assess within-individual associations: that is, whether the same person is capable of accurately track day-to-day fluctuations in the quality his/her sleep via self-reports.

Within-individual associations are free from trait-level confounding as the same persons’ characteristics influencing both objective sleep quality and subjective sleep quality ratings (such as age or personality) do not meaningfully change over the course of a sleep study, typically lasting one or two weeks at most. Significant within-individual associations between objective and subjective sleep quality – for example, if the same person reports better sleep in the morning after nights with higher sleep efficiency, both variables compared to his/her average – are strong evidence for the correspondence of objective and subjective metrics.

Recently, Shirota et al (Shirota et al., 2023) performed a study off 77 health adults undergoing polysomnography and reporting sleep quality the subsequent morning. The study used an innovative design in which all participants spent two nights in the laboratory, and differences between the two nights in objective and subjective sleep quality were compared, essentially performing a within-participant analysis. In this study, increases in total quality sleep were significantly correlated with increases in time spent in N3 sleep and delta EEG power, while increases in subjectively rated depth of sleep were correlated with increases in sleep efficiency, time spent in N3 sleep, delta EEG power, and decreases in WASO (|r|=0.405-0.438). However, this study was limited by the low number of nights sampled and the basic statistical approach. In another within-individual study (Svetnik et al., 2020), compared subjective ratings on sleep across multiple nights in insomnia patients undergoing drug treatment and found sthat total sleep time correlates the best with subjective sleep ratings. This study is, however, limited by its use of pharmacological treatment (known to patients in their crossover design) to influence sleep, the lack of sleep efficiency as a predictor, and the fact that repeated sleep measurements were often weeks or months apart. We are unaware of further studies with a within-participant design. The scoping review by McCarter et al (McCarter et al., 2022) also does not mention any multiday observational study.

Therefore, our aim with the current study was to bridge this gap and perform the first multiday observational study of subjective and objective sleep quality. We used the Budapest Study of Traits, Experiences and Sleep (BSETS), a new large dataset of over 250 participants with seven consecutive nights of mobile EEG data and subjective sleep ratings to calculate how subjective and objective sleep quality correlates between individuals, as well as between the same nights of the same individual.

## Methods

### Participants

We used data from the Budapest Sleep, Experiences and Traits Study (BSETS). The full protocol of this dataset, including a description of available data, has been published separately (Taji et al., 2023). In brief, BSETS is a multiday observational study in which healthy volunteering participants fill out diaries each evening and each morning for seven consecutive days and record their sleep with a Dreem2 mobile EEG headband (Dreem Inc, 2017).

The Institutional Review Board (IRB) of Semmelweis University as well as the Hungarian Medical Council (under 7040-7/2021/ EÜIG “Vonások és napi események hatása az alvási EEG-re” (The effect of traits and daily activities and experiences on the sleep EEG)) approved BSETS as compliant with the latest revision of the Declaration of Helsinki. All participants gave written informed consent on a form reviewed and approved by the IRB.

Full observations (hypnograms, electrophysiology and questionnaire data, including lagged outcomes from the previous nights which rendered the first night of each participant unusable) were available from 1318 nights, recorded from 246 participants. Some additional data loss was observed in models with more variables (see Results for detailed sample sizes).

### Objective sleep quality

Participant slept on each night with the Dreem2 mobile EEG headband device which recorded quantitative EEG. Recordings were automatically scored with an algorithm that demonstrated high validity against visual scorings (Arnal et al., 2020). Objective sleep ratings were extracted from the hypnogram created in this way. Objective sleep ratings used in the current study were the following: sleep efficiency (SE), total sleep time (TST), sleep onset latency (SOL), wake after sleep onset (WASO), N2 latency, N3 latency, REM latency, and the number of awakenings.

### Quantitative EEG analysis

Based on previous analyses of quantitative EEG recorded with the Dreem2 headband (Taji et al., 2023) we chose the channel F7-O1 for EEG analyses due to a good compromise of data quality/availability and large electrode distance allowing the recording of topographically widespread activity such as slow waves. Data was recorded using dry silicone electrodes and a sampling frequency of 250 Hz (see (Taji et al., 2023) and (Dreem Inc, 2017) for technical details). A complimentary algorithm scored the quality of EEG data segments on a 2-second basis. Data was discarded if this algorithm gave an artifact probability of greater than 25%. If channel quality (the proportion of data epochs from that channel with lower than 25% artifact probability) was 20% for a night, data from this channel was discarded. These settings were based on preliminary analyses (Taji et al., 2023) suggesting that these settings result in the best tradeoff off data quality and data availability.

We used the periodogram() function in MATLAB EEGLab with 2-second nonoverlapping epochs and Hamming windows to perform spectral analysis. Power spectral density (PSD) estimates were averaged for each night. We used PSD data from the low sigma frequency band (10-13 Hz) in N2 sleep to estimate sleep spindling, and the delta frequency band (0.5-4 Hz) in SWS to estimate slow wave activity. The use of the slow rather than the fast sigma frequency range was motivated by the fact that the frontal electrode setup of Dreem2 results in sleep spindle peaks in this range (Taji et al., 2023). PSD estimates were log-transformed and relativized before analysis.

### Subjective sleep quality

Upon awakening, participants filled out the Groningen Sleep Quality Scale (GSQS) (Simor et al., 2009), a 15-item scale (with 1 unscored item) in which participants subjectively rate the quality of their sleep with a set of yes/no questions. Our primary outcome of interest was GSQS total score. Higher scores indicate lower sleep quality.

For sensitivity analyses and to replicate our original findings, we considered two additional outcomes of interest: 1) an additional question in which participants are asked to rate their level of restedness from 1 to 10 on a Likert scale, considered as a continuous variable and 2) response to the first unscored question of the GSQS (“I had a deep sleep last night”), a binary variable. On both alternative outcomes of interest, a higher score indicates higher sleep quality.

### Statistical analysis

For each predictor, we created two versions of the original variable (McCrae et al., 2008; Robson and Pevalin, 2015). The first contained contained within-individual differences (defined as the original value minus the mean of the individual). This variable was used to estimate Level 1 (within-individual) effects. The second contained the individual means. This variable was used to estimate Level 2 (between-individual) effects.

In initial analyses, we calculated simple Pearson correlations between Level 1 (within-individual correlations) and Level 2 (between-individual correlations) variables. Within-individual correlations express whether deviations from the individual mean on two variables are correlated, for example, if the same person reports better than average sleep after nights with more than average N3 sleep. Between-individual correlations express whether the typical values of participants resemble, for example, whether participants usually reporting better sleep also usually experience more N3 sleep. These correlations, however, are not free from confounders. For example, within-individual correlations may be confounded by day of week, and between-individual correlations may be confounded by age. This would mean that on weekends, participants experience deeper sleep and report higher satisfaction with sleep without an actual causal link, while younger participants typically have deeper sleep and higher satisfaction with sleep, again without an actual causal link.

In subsequent analyses, we used multilevel models implemented in MATLAB using the fitglme() function. to simultaneously estimate Level 1 and Level 2 effects and control for confounders. All models were controlled for age, sex at Level 2 and day of the week (weekday/weekend, defined as a binary variable) and the previous mornings’ GSQS score at Level 1. This control for lagged outcomes is necessary to eliminate the effects of recovery sleep after nights of poor sleep, after which deeper sleep and improved sleep subjective quality are logically expected. A random intercept by participant was added to each model.

We set up a series of increasingly inclusive models to estimate the effects of objective sleep metrics on subjective sleep quality on an increasingly exploratory basis.

First, we fitted a baseline model with only the control variables (day of the week, lagged subjective sleep quality, age, sex and a random intercept by participant) as predictors. These models served to evaluate the amount of variance accounted for by variables of no interest, to compare them to models also including objective sleep metrics.

In Model 1, we only used sleep efficiency as a predictor, as this is the most inclusive single objective metric of sleep quality with reasonably high correlations with more refined other metrics (**Table 1**).

**Table 1.** Descriptive statistics and correlations of the variables in the analysis. For correlations, day of the week and sex were coded as binary variables with 1 standing for ‘weekend’ and ‘male’, respectively, so positive correlations with these variables mean higher values in these categories and the mean value refers to their proportion in the sample.

|  | Mean | SD | Groningen<br>Total<br>Score | Sleep<br>Efficiency | SOL | WASO | TST | N2 % | N3 % | REM % | N2<br>Latency | N3<br>Latency | REM<br>Latency | Awake<br>nings | Sigma<br>Power | Delta<br>Power | Weekend | Age |
| --- | --- | --- | --- | --- | --- | --- | --- | --- | --- | --- | --- | --- | --- | --- | --- | --- | --- | --- |
| Groningen<br>Total Score | 4,128 | 3,408 |  |  |  |  |  |  |  |  |  |  |  |  |  |  |  |  |
| Sleep<br>Efficiency | 91,060 | 6,698 | -0.402 |  |  |  |  |  |  |  |  |  |  |  |  |  |  |  |
| SOL | 14,849 | 14,702 | 0,218 | -0.633 |  |  |  |  |  |  |  |  |  |  |  |  |  |  |
| WASO | 20,964 | 21,485 | 0,275 | -0.668 | 0,150 |  |  |  |  |  |  |  |  |  |  |  |  |  |
| TST | 396,53 | 91,742 | -0.285 | 0,341 | -0,099 | 0,084 |  |  |  |  |  |  |  |  |  |  |  |  |
| N2 % | 46,297 | 9,176 | 0,019 | -0.029 | 0,038 | 0,192 | 0,302 |  |  |  |  |  |  |  |  |  |  |  |
| N3 % | 21,827 | 9,526 | -0.007 | 0,142 | -0,097 | -0,313 | -0,407 | -0,669 |  |  |  |  |  |  |  |  |  |  |
| REM % | 25,796 | 7,507 | -0.097 | -0,014 | 0,041 | <0,001 | 0,161 | -0,439 | -0,333 |  |  |  |  |  |  |  |  |  |
| N2 Latency | 4,350 | 4,953 | 0,076 | -0,102 | 0,140 | 0,071 | 0,001 | -0,014 | -0,028 | -0,009 |  |  |  |  |  |  |  |  |
| N3 Latency | 18,522 | 18,890 | 0,145 | -0.246 | 0,095 | 0,272 | -0,059 | 0,220 | -0,259 | -0,020 | 0,246 |  |  |  |  |  |  |  |
| REM<br>Latency | 79,347 | 36,839 | 0,048 | <0,001 | -0,027 | 0,071 | 0,075 | 0,134 | 0,102 | -0,310 | 0,011 | -0,054 |  |  |  |  |  |  |
| Awakenings | 19,542 | 9,487 | 0,025 | -0.192 | 0,076 | 0,516 | 0,469 | 0,237 | -0,451 | 0,087 | 0,043 | 0,081 | 0,067 |  |  |  |  |  |
| Sigma<br>Power | -0,169 | 0,137 | 0,042 | -0.007 | -0,057 | -0,017 | -0,119 | -0,005 | 0,045 | -0,008 | -0,047 | -0,013 | <0,001 | -0,088 |  |  |  |  |
| Delta Power | 1,467 | 0,189 | -0.030 | 0,120 | -0,053 | -0,158 | -0,077 | -0,305 | 0,420 | -0,083 | 0,023 | -0,115 | 0,097 | -0,169 | 0,374 |  |  |  |
| Weekend | 0,283 | 0,451 | -0.057 | 0,020 | -0,023 | 0,027 | 0,024 | 0,019 | -0,037 | 0,022 | 0,022 | 0,026 | -0,051 | 0,016 | -0,004 | -0,003 |  |  |
| Age | 28,810 | 12,672 | 0,075 | -0.195 | 0,005 | 0,274 | -0,011 | 0,320 | -0,370 | -0,008 | -0,081 | 0,124 | -0,095 | 0,120 | -0,107 | -0,556 | -0,002 |  |
| Sex | 0,552 | 0,497 | 0,033 | -0,018 | 0,039 | -0,002 | 0,060 | 0,049 | -0,052 | 0,057 | -0,025 | -0,001 | -0,036 | -0,094 | 0,245 | 0,214 | 0,000 | 0,013 |

In Model 2, we added the predictors sleep onset latency, wake after sleep onset and total sleep time. These metrics are more specific than sleep efficiency and have been linked to subjective sleep quality in previous studies (see Introduction). We removed sleep efficiency from Model 2 and from subsequent models due to multicollinearity (see high correlations with other metrics in **Table 1**).

In Model 3, we added further objective sleep metrics to explore the role of more refined sleep composition on sleep quality. These were the percentage and latency of N2, N3 and REM and the number of awakenings.

Finally, in Model 4, we added two quantitative EEG metrics, N2 low sigma relative power and N3 delta relative power, as predictors. These were included following a previous study (Gabryelska et al., 2019) reporting a correlation between similar metrics and subjective sleep quality. Low sigma was used because the frontal channels in BSETS map slow spindles in this frequency range better (Taji et al., 2023). Both metrics were derived from the channel F7-O1 which demonstrated favorable characteristics in preliminary analyses (Taji et al., 2023).

Raw data and code to replicate analyses are available at: https://zenodo.org/records/10427245.

## Results

### Descriptive statistics

Descriptive statistics (means, standard deviations and a correlation matrix) are reported in **Table 1**.

### Within-and between-participant correlations

As an initial step of our analyses, we calculated intraclass correlation coefficients (variance accounted for by participant ID), between-person correlations (correlations of individual means), and within-person correlations (correlations of deviations from individual means). Findings are reported in **Table 2**.

**Table 2.**
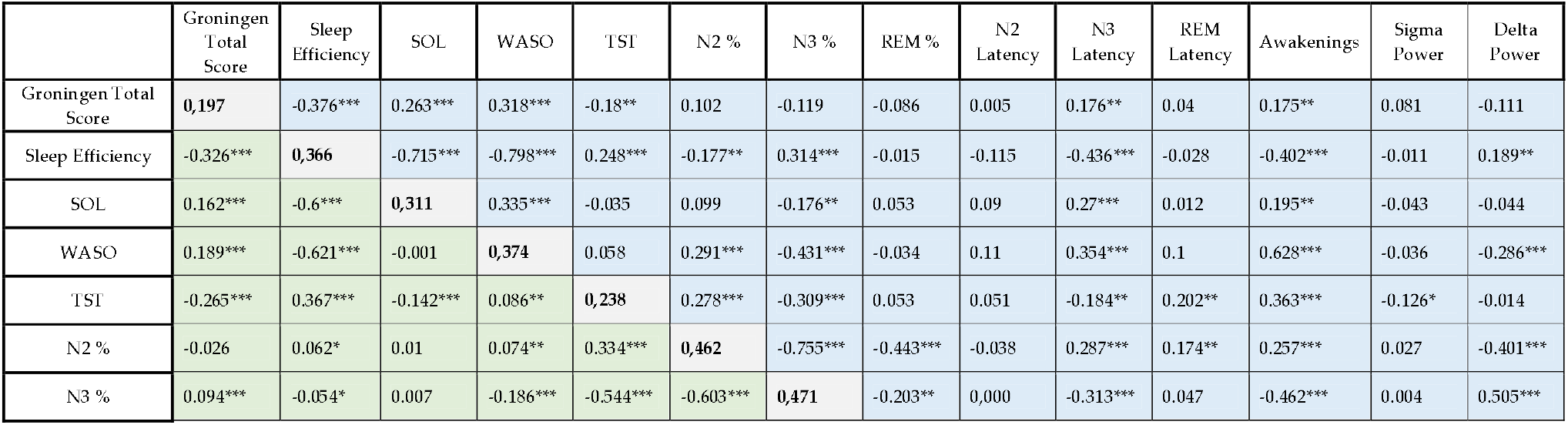
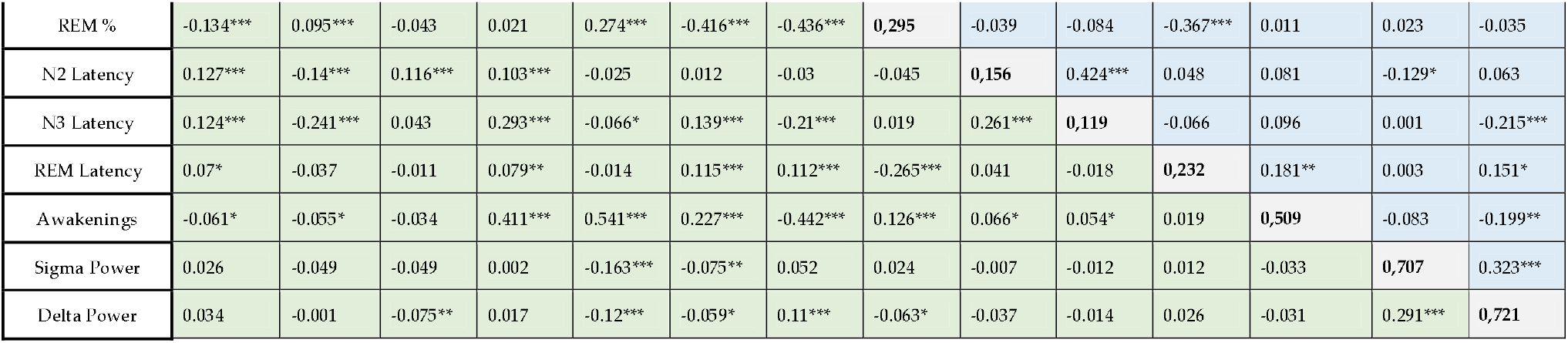
Within- and between-person similarity of variables. Values in the diagonal (grey) show intraclass correlation coefficients (variance accounted for by participant ID) in bold. Values above the diagonal (blue) are between-person correlation values (correlations of individual means). Values below the diagonal (green) are within-person correlations (correlations of deviations from the individual mean). (p<0.05*, p<0.01**, p<0.001***)

We found high intraclass coefficients (high within-individual similarity) of EEG PSD values. For the other variables, intraclass correlation coefficients were moderate to substantial. The lowest value was found for N3 latency (ICC=0.119), and the highest for the number of awakenings (ICC=0.509).

GSQS total scores exhibited significant between-participant correlations with sleep onset latency (SOL), wake after sleep onset (WASO) (r=0.318, p<0.001) and total sleep time (TST). Within-participant correlations were similarly strong (|r|=0.124-0.326) and due to the much larger number of nights than participants, strongly significant (p<0.0001 in all cases).

### Multilevel modelling

In Model 1, we found that sleep efficiency was significantly related to sleep quality both at the between- and the within-individual levels. Linear estimates suggested that participants with a one percent higher mean sleep efficiency can expect a 0.17 point lower average score on the GSQS. Within-participant estimates were even higher, with a 0.25 point drop in scores expected for each additional percent of sleep efficiency relative to the individual mean. 16% of subjective sleep quality variance was accounted for by sleep efficiency alone over the null model.

In Model 2, we replaced sleep efficiency with total sleep time, sleep onset latency and wake after sleep onset. All were related to significantly related to subjective sleep quality both at the within- and between-individual levels. Again, within-individual estimates were ∼20-100% higher, suggesting that studying within-individual differences is a better way to capture the relationship between objective and subjective sleep quality. The three objective sleep metrics together accounted for 19% of the variance of self-reported sleep quality.

In Model 3, additional sleep macrostructure metrics were added. Higher N2, N3 and REM percentage were all significantly related to better sleep quality, however, only at the within-individual level. N2, N3 and REM latency, and the number of awakenings were unrelated to subjective sleep quality, except for a borderline significant within-individual estimate for N2 latency (p=0.046). Estimates for the previously entered metrics did not substantially change, suggesting that different components of the sleep macrostructure have largely independent associations with sleep quality. Despite the significant effects, variance accounted for improved only marginally by 1%.

In Model 4, qEEG metrics N2 low sigma and N3 delta relative power were added. These were not associated with sleep quality either at the between- or within-individual levels. The model including EEG metrics only accounted for 17% of subjective sleep rating variance. However, this model is not directly comparable to previous ones because of missingness in the EEG data affecting the amount of available data.

**Table 3** summarizes within-individual effects across the four models. **Figure 1** illustrates within-participant correlations.

**Table 3.** Within-participant effects on subjective sleep quality. The table contains fixed effects associated with the deviation of subjective sleep quality metrics from individual means. The table contains unstandardized regression coefficients, showing the expected change in GSQS points as a function of a one-unit increase in sleep metrics. Sleep metrics are expressed as percentage points for sleep efficiency and sleep composition, minutes for total sleep time and sleep latency, total number for awakenings and log10 microvolt/sec^2^ for relative power. Incremental R^2^ refers to the variance accounted for by the models in addition to the variance accounted for by the random intercept and control variables. R^2^ values are shown for the full model, not only within-individual effects.

|  | Model 1 |  |  | Model 2 |  |  | Model 3 |  |  | Model 4 |  |  |
| --- | --- | --- | --- | --- | --- | --- | --- | --- | --- | --- | --- | --- |
|  | B | SE | p | B | SE | p | B | SE | p | B | SE | p |
| Sleep Efficiency | -0,251 | 0,017 | <0.001 |  |  |  |  |  |  |  |  |  |
| WASO |  |  |  | 0,047 | 0,005 | <0.001 | 0,040 | 0,006 | <0.001 | 0,041 | 0,006 | <0.001 |
| TST |  |  |  | -0,013 | 0,001 | <0.001 | -0,012 | 0,002 | <0.001 | -0,012 | 0,002 | <0.001 |
| SOL |  |  |  | 0,032 | 0,007 | <0.001 | 0,029 | 0,007 | <0.001 | 0,032 | 0,008 | <0.001 |
| REM % |  |  |  |  |  |  | -0,215 | 0,062 | 0,001 | -0,213 | 0,069 | 0,002 |
| REM Latency |  |  |  |  |  |  | 0,005 | 0,003 | 0,075 | 0,004 | 0,003 | 0,141 |
| N3 % |  |  |  |  |  |  | -0,232 | 0,063 | <0.001 | -0,228 | 0,071 | 0,001 |
| N3 Latency |  |  |  |  |  |  | 0,005 | 0,005 | 0,280 | 0,005 | 0,007 | 0,449 |
| N2 % |  |  |  |  |  |  | -0,199 | 0,064 | 0,002 | -0,194 | 0,071 | 0,007 |
| N2 Latency |  |  |  |  |  |  | 0,035 | 0,018 | 0,046 | 0,078 | 0,022 | <0.001 |
| Awakenings |  |  |  |  |  |  | -0,033 | 0,019 | 0,077 | -0,035 | 0,021 | 0,099 |
| Delta Power |  |  |  |  |  |  |  |  |  | 0,892 | 0,935 | 0,340 |
| Sigma Power |  |  |  |  |  |  |  |  |  | 0,088 | 1,237 | 0,943 |
|  | Number of participants |  | 246 | Number of participants |  | 246 | Number of participants |  | 246 | Number of participants |  | 235 |
|  | Number of nights |  | 1318 | Number of nights |  | 1318 | Number of nights |  | 1308 | Number of nights |  | 1092 |
|  | Incremental R-squared |  | 0,16 | Incremental R-squared |  | 0,19 | Incremental R-squared |  | 0,20 | Incremental R-squared |  | 0,17 |

**Figure 1.**
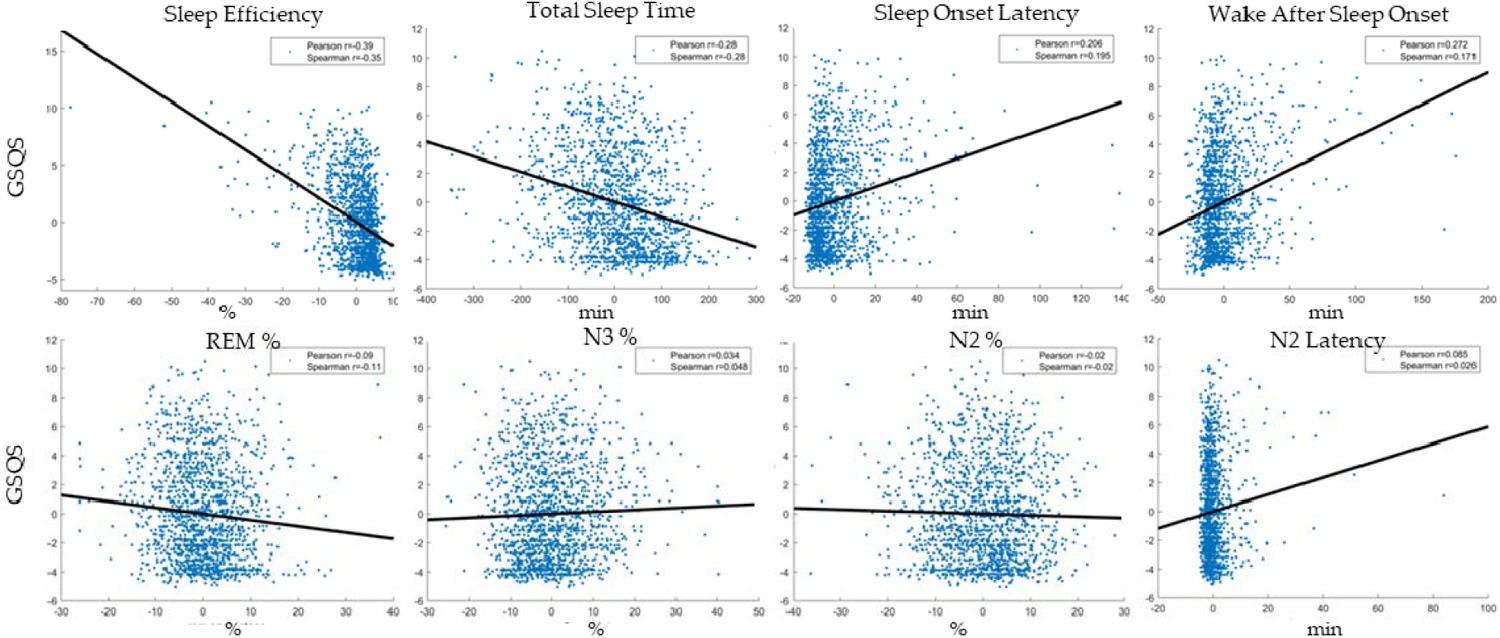
Within-participant associations between indicators of subjective sleep quality (GSQS total score, vertical axis) and objective sleep metrics (separate panels, horizontal axis). The scatterplots show deviations from the individual means, pooled across participants. In order to illustrate partial correlations net of confounders, control variables (age, sex, day of week, lagged outcomes) were regressed out of deviations. Because the plots show raw residuals, data points are centered around 0. Sleep metrics are selectively shown if they reached nominal significance (p<0.05) in Model 4 (**Table 3**). Both Pearson and Spearman correlations are shown to illustrate the effect of outliers on the associations.

**Table 4** summarizes between-individual effects. **Figure 2** illustrates between-individual correlations.

**Table 4.** Between-participant effects on subjective sleep quality. The table contains fixed effects associated with the individual means of objective sleep quality, regressed on mean GSQS scores. The table contains unstandardized regression coefficients, showing the expected change in GSQS points as a function of a one-unit increase in sleep metrics. Sleep metrics are expressed as percentage points for sleep efficiency and sleep composition, minutes for total sleep time and sleep latency, total number for awakenings and log10 microvolt/sec^2^ for relative power.

|  | Model 1 |  |  | Model 2 |  |  | Model 3 |  |  | Model 4 |  |  |
| --- | --- | --- | --- | --- | --- | --- | --- | --- | --- | --- | --- | --- |
|  | B | SE | p | B | SE | p | B | SE | p | B | SE | p |
| Sleep Efficiency | -0,171 | 0,025 | <0,001 |  |  |  |  |  |  |  |  |  |
| TST |  |  |  | 0,042 | 0,008 | <0,001 | 0,040 | 0,011 | <0,001 | 0,032 | 0,013 | 0,013 |
| SOL |  |  |  | -0,006 | 0,002 | 0,001 | -0,006 | 0,003 | 0,022 | -0,005 | 0,003 | 0,106 |
| WASO |  |  |  | 0,026 | 0,012 | 0,036 | 0,024 | 0,013 | 0,057 | 0,033 | 0,013 | 0,011 |
| REM % |  |  |  |  |  |  | -0,059 | 0,105 | 0,571 | -0,167 | 0,113 | 0,140 |
| REM Latency |  |  |  |  |  |  | 0,000 | 0,005 | 0,965 | -0,003 | 0,006 | 0,639 |
| N3 % |  |  |  |  |  |  | -0,056 | 0,103 | 0,589 | -0,136 | 0,111 | 0,219 |
| N3 Latency |  |  |  |  |  |  | 0,003 | 0,015 | 0,849 | -0,002 | 0,016 | 0,915 |
| N2 % |  |  |  |  |  |  | -0,059 | 0,107 | 0,584 | -0,159 | 0,115 | 0,165 |
| N2 Latency |  |  |  |  |  |  | -0,036 | 0,049 | 0,457 | -0,014 | 0,050 | 0,780 |
| Awakenings |  |  |  |  |  |  | -0,005 | 0,030 | 0,860 | -0,024 | 0,032 | 0,457 |
| Delta Power |  |  |  |  |  |  |  |  |  | -1,645 | 1,017 | 0,106 |
| Sigma Power |  |  |  |  |  |  |  |  |  | 1,806 | 1,042 | 0,083 |

**Figure 2.**
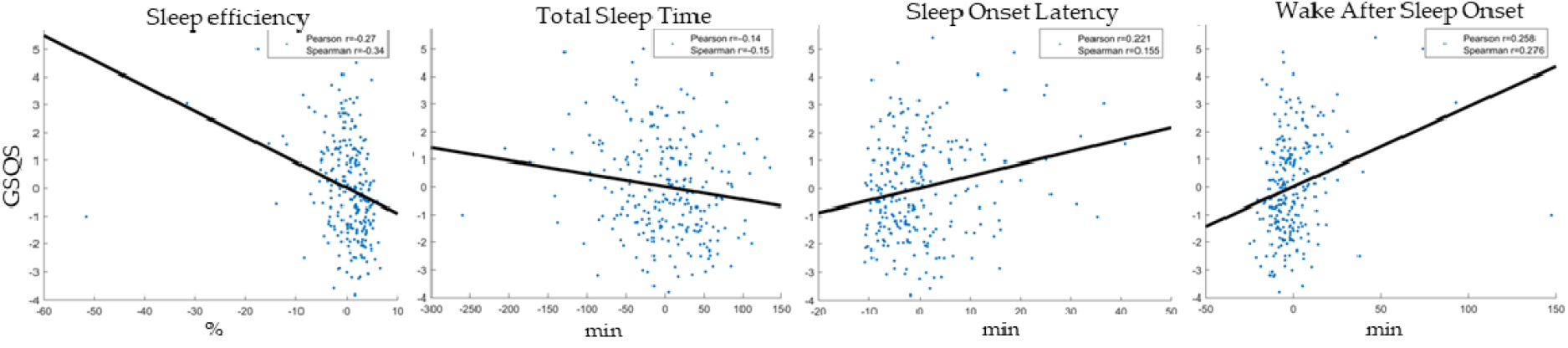
Between-participant associations between indicators of subjective sleep quality (GSQS total score, vertical axis) and objective sleep metrics (separate panels, horizontal axis). The scatterplots show individual means. In order to illustrate partial correlations net of confounders, control variables (age, sex, day of week, lagged outcomes) were regressed out of deviations. Because the plots show raw residuals, data points are centered around 0. Sleep metrics are selectively shown if they reached nominal significance (p<0.05) in Model 4 (**Table 4**). Both Pearson and Spearman correlations are shown to illustrate the effect of outliers on the associations.

### Sensitivity analyses

In order to further confirm our findings, we re-ran analyses using two alternative operationalizations of subjective sleep quality: 1) the first question of the GSQS (“I had a deep sleep last night”), which is not counted towards the total score, 2) a custom question prompting participants to rate their level of restedness on a Likert scale from 1 to 10. These variables were moderately correlated with GSQS total score (r=-0.64 and r=0.25, respectively) and with each other (r=-0.16).

For the first GSQS question, a generalized mixed model with a logit link function was fitted, using the fitglme() MATLAB function. Results were replicated, with significant within-participant effects of sleep efficiency, total sleep time, sleep onset latency, wake after sleep onset, REM latency, N2, N3 and REM percentage. Between-participant effects were again weaker, and only significant for sleep efficiency and wake after sleep onset. For example, for each additional percentage of sleep efficiency, the within-participant odds ratio for participants reporting having had “a deep sleep last night” was 1.13, and for each additional minute of wake after sleep onset it was 0.98.

Self-reported morning restedness also exhibited significant within-participant relationships with sleep efficiency, wake after sleep onset and total sleep time (but not sleep onset latency). For this variable, no other significant within-participant and no between-participant effects were found.

Detailed statistics about alternative subjective sleep quality ratings are reported in **Supplementary Tables S1-S4**.

### Terminal sleep stages

In a final additional analysis, we investigated if terminal sleep stages (the sleep stage last experienced by participants before awakening) affects perceived sleep quality. This analysis was motivated by recent research (Stephan et al. 2021) using an awakening protocol which found that perceived sleep depth differs as a function of the type of sleep participants are awakened from. We defined the terminal sleep stage as the last sleep stage scored before the final awakening at the end of recordings. The terminal sleep stage was added as a predictor to Model 3. Compared to the reference category, waking up from N2 sleep was associated with a trend of higher subjective sleep quality amounting to about 1/3 of a GSQS point (B=-0.354, p=0.047), but no effect was found for N1 or SWS.

Because we were concerned about scoring fidelity issues when only using the very last epoch as the terminal sleep stage, we investigated two stricter model specifications. In the first, terminal sleep stage was defined as the mode of the last 10 scored epochs. In the second, even stricter specification, the mode was only used if the frequency of the mode was at least 50% of the last 10 epochs, and otherwise data was set to missing. Both specifications confirmed higher subjective sleep quality after waking from N2 (B=-0.496, p=0.003; and B=-0.523, p=0.002, respectively) than REM, but none from the other sleep stages. Incremental R^2^ values (adjusted for degrees of freedom) for these models did not exceed that of Model 3.

In sum, while awakening from N2 is associated with higher subjective sleep quality ratings compared to other sleep stages, but terminal sleep stage accounts for a negligible amount of variance in GSQS scores.

## Discussion

In our work, using a large sample of health volunteers undergoing at-home EEG monitoring for a full week, we showed clear evidence that subjective sleep ratings are moderately related to objective sleep metrics. The same participant tended to report better sleep after nights where sleep was, based on EEG-based measures, objectively better. Sleep efficiency was the most important variable predicting subjective sleep quality, but many other variables including faster sleep latency, increased time in N2, N3 and REM, as well as reduced WASO also contributed. We found no evidence that sleep EEG power or the number of awakenings independently contribute to better subjective sleep ratings. Our findings were robust to the choice of subjective sleep ratings. Overall, objective sleep metrics only accounted for about one fifth of subjective ratings of sleep.

Our findings contradict the conclusions of previous review (McCarter et al., 2022) as well as several large-sample papers (Kaplan et al., 2017b, 2017a) that subjective sleep quality is at best weakly related to objective sleep metrics. Our findings, in line with studies using a similar within-participant methodology (Shirota et al., 2023; Svetnik et al., 2020), indicate that up to 20% of the variance in subjective sleep ratings can be accounted for by objective sleep metrics. This discrepancy likely arises from methodological issues, specifically the assessment of habitual or current sleep quality as the subjective indicator, univariate or multivariate models, and the use of between-participant or within-participant designs. Our view is that within-participant designs using current subjective sleep quality are the only ones which can give an unbiased estimated of the concordance of subjective and objective sleep ratings.

First, it is questionable if subjective estimates of habitual sleep quality are expected to reflect objective sleep metrics on a given night. One study (Unruh et al., 2008) used a large sample (the MrOS Sleep Study) and discovered no association between subjectively assessed habitual sleep quality and polysomnography-assessed sleep on a single night. A similar analysis of an overlapping dataset using current sleep quality assessed after the laboratory night, conversely, found significant associations (Kaplan et al., 2017b). Self-reports of habitual sleep quality may have dubious accuracy and may be contaminated by personality variables or biased responding styles, such as neuroticism, mood or participants’ preference for disclosing medical issues. Conversely, a single measure of objective sleep (even if an adaptation night or an average of several nights is used) may be also be an imperfect estimate of how an individual typically sleeps. Thus, negative findings from studies using habitual self-reports of sleep quality are not a strong argument against a link between objective and subjective sleep quality.

Second, even if current sleep quality is used, subjective sleep assessment may be biased by personality or responding styles. For example, individuals more prone to depression or those with a greater willingness to talk about poor health may report worse sleep even given the same objective sleep experience, biasing subjective-objective correlations if a single estimate of each is used. Because response tendencies introduce noise, the expected direction of the bias is downward, underestimating subjective-objective correlations. Such biases may contribute to the frequent absence of such associations (McCarter et al., 2022) in the previous literature. Ideally, the same individuals would be followed up for several days and correlated fluctuations in objective and subjective sleep quality would serve as evidence for the relationship of the two. Two recent studies (Shirota et al., 2023; Svetnik et al., 2020) employing such a design indeed found a substantial association between subjective and objective sleep quality.

In our analyses, between-participant associations were found between sleep efficiency, sleep latency, WASO, total sleep time and subjective sleep quality. In other words, participants usually having a higher ratio of bedtime spent asleep also typically reported a better subjective experience of sleep, even controlling for age and sex. These findings are analogous to between-participant studies reporting objective-subjective associations. One way to interpret between-participant effects is that participants’ subjective reports are accurate reflections of their better sleep. This is indeed the interpretation endorsed by most previous studies with this design. However, average total scores on the GSQS across the seven nights are also correlated with scores on the Emotional instability subscale of the Big Five Inventory personality scale (r=0.278, p=10^-6^), the Neuroticism-Anxiety subscale of the Zuckerman-Kuhlman Personality Questionnaire (r=0.245, p=10^-4^), weekly average negative emotional ratings of their days on the Positive and Negative Affect Scale (r=0.274, p=10^-6^), and the mean subjective ratings of their days as “Happy” as opposed to “Sad” on a Likert scale (r=-0.189, p=0.002). (See the BSETS protocol paper (Taji et al., 2023) for details on these variables.) Thus, an alternative explanation is that GSQS scores partially reflect the tendency, related to trait neuroticism, of some individuals to see life experiences – including their sleep – in a negative light. While individuals with higher levels of this trait typically experience worse sleep, the correlation with subjective ratings may be accidental and does not necessarily reflect an accurate perception of reduced sleep quality.

Crucially for a causal interpretation we found that the associations between subjective and objective sleep quality persist in within-individual analyses. After a night with objectively worse sleep quality (indicated by reduced sleep efficiency, total sleep time or the percentage of N2 or REM, or by increased WASO, sleep latency, but also of N3 sleep percentage) the same participant tended to report worse subjective sleep quality as well. These findings are based on time-lagged measures from two independent data sources (EEG machine and subjective experience) and due to the within-participant nature of analyses cannot be biased by trait-level confounders such as personality. Therefore a causal interpretation is warranted: participants can accurately perceive on which night their sleep was objectively better. Objective sleep parameters accounted for close to 20% of the variance of subjective sleep ratings (over the effect of controls).

A similar conclusion was reached by a large clinical study (Svetnik et al., 2020) which found that 14-27% of subjective sleep ratings could be accounted for by objective sleep metrics and controls such as age. This study, however, did not calculate sleep efficiency which we found to be the single strongest correlate of subjective sleep ratings. It was also based on data from a drug trial where changes in sleep quality were drug-induced and participants may have been aware of this fact. Another study (Shirota et al., 2023) also found substantial within-participant correlation with, for instance, sleep efficiency accounting for 9% of overall subjective sleep quality and 16% of the perceived depth of sleep. (Effect size conversions from the original AUC estimates and correlations were performed using www.escalc.site). Ours, however, is the first study to rigorously model both between-participant and within-individual effects of a comprehensive set of objective sleep metrics in a large sample of healthy volunteers to demonstrate that these are related to the subjective perception of sleep.

The main limitation of our study is that objective sleep metrics were not obtained from polysomnography but via an ambulatory EEG device. While automatically scored hypnograms from this device are very similar to expert ratings (Arnal et al., 2020) and we also confirmed the validity of EEG characteristics (Taji et al., 2023), imperfections in the measurement of objective sleep may have attenuated its association with subjective assessments of it. Furthermore, as our sample mostly consisted of health young volunteers, the results may generalize less perfectly to other populations. While our findings confirmed that objective and subjective sleep ratings are related, only a modest amount of variance is accounted for, leaving the main sources of subjective sleep quality experiences undiscovered.

Our findings have implications for both health care and research. As sleep problems are common as a significant burden (Grandner, 2022), interventions improving subjective sleep have paramount importance. Our research shows indicates that the greatest improvement in subjective sleep quality can be expected if overall sleep efficiency is improved, while other intervention targets (e.g. changing sleep composition via medication or introducing specific EEG oscillations via stimulation or neurofeedback) are less promising. Nevertheless, we found that the association between objective and subjective sleep quality is weak, warranting caution about the expected effect of interventions. Given the modest strength of this association, even great improvements in objective sleep quality will likely only result in a modestly improved sleep experience. Researchers must also be aware that subjective sleep quality is relatively weakly related to objective sleep metrics, thus, they cannot be treated as analogs or proxy measures of each other in scientific studies.

## CRediT Author Statement

RPB: Investigation, Visualization Data Curation, Writing – Original Draft, Writing – Review and Editing.

PPU: Conceptualization, Methodology, Funding acquisition, Project administration, Supervision, Software, Formal Analysis, Writing – Original Draft, Writing – Review and Editing.

## Acknowledgements

This was supported by the National Research, Development and Innovation Office – NKFIH (grant number: 138935).

