## Supplementary table for "Objective sleep quality predicts subjective sleep ratings: a multiday observational study"

Supplementary tables

|  | Model 1 | | Model 2 | | Model 3 | | Model 4 | |
| --- | --- | --- | --- | --- | --- | --- | --- | --- |
|  | OR | p | OR | p | OR | p | OR | p |
| Sleep Efficiency | 1,132 | <0.001 |  |  |  |  |  |  |
| TST |  |  | 0,977 | <0.001 | 0,985 | 0,004 | 0,979 | 0,001 |
| SOL |  |  | 1,004 | <0.001 | 1,005 | <0.001 | 1,005 | 0,004 |
| WASO |  |  | 0,984 | 0,006 | 0,985 | 0,020 | 0,983 | 0,016 |
| REM % |  |  |  |  | 1,156 | 0,012 | 1,157 | 0,028 |
| REM Latency |  |  |  |  | 0,995 | 0,030 | 0,994 | 0,034 |
| N3 % |  |  |  |  | 1,223 | 0,001 | 1,219 | 0,004 |
| N3 Latency |  |  |  |  | 1,003 | 0,615 | 1,007 | 0,287 |
| N2 % |  |  |  |  | 1,162 | 0,011 | 1,144 | 0,049 |
| N2 Latency |  |  |  |  | 0,990 | 0,553 | 0,965 | 0,072 |
| Awakenings |  |  |  |  | 0,991 | 0,588 | 0,989 | 0,598 |
| Delta Power |  |  |  |  |  |  | 0,513 | 0,486 |
| Sigma Power |  |  |  |  |  |  | 2,676 | 0,444 |
|  | Number of participants | 246 | Number of participants | 246 | Number of participants | 246 | Number of participants | 235 |
|  | Number of nights | 1375 | Number of nights | 1375 | Number of nights | 1364 | Number of nights | 1141 |
|  | Incremental R-squared | 0,33 | Incremental R-squared | 0,11 | Incremental R-squared | 0,11 | Incremental R-squared | 0,13 |

**Supplementary Table S1.** Within-participant effects on subjectively rated deep sleep. The table contains fixed effects associated with the deviation of objective sleep quality metrics from individual means. The table presents odds ratios, indicating the impact of a one-unit change in objective sleep quality metrics on the odds of a response indicating deep sleep based on first GSQS item. Sleep metrics are expressed as percentage points for sleep efficiency and sleep composition, minutes for total sleep time and sleep latency, total number for awakenings and log10 microvolt/sec^2^ for relative power. Incremental R^2^ refers to the variance accounted for by the models in addition to the variance accounted for by the random intercept and control variables. R^2^ values are shown for the full model, not only within-individual effects.

|  |  | Model 1 |  |  | Model 2 |  |  | Model 3 |  |  | Model 4 |  |
| --- | --- | --- | --- | --- | --- | --- | --- | --- | --- | --- | --- | --- |
|  | B | SE | p | B | SE | p | B | SE | p | B | SE | p |
| Sleep Efficiency | -0,051 | 0,012 | <0.001 |  |  |  |  |  |  |  |  |  |
| TST |  |  |  | 0,009 | 0,004 | 0,013 | 0,009 | 0,004 | 0,031 | 0,009 | 0,005 | 0,073 |
| SOL |  |  |  | -0,004 | 0,001 | <0.001 | -0,003 | 0,001 | 0,009 | -0,004 | 0,001 | 0,004 |
| WASO |  |  |  | 0,005 | 0,005 | 0,340 | 0,005 | 0,005 | 0,367 | 0,008 | 0,006 | 0,196 |
| REM % |  |  |  |  |  |  | -0,025 | 0,047 | 0,599 | -0,002 | 0,052 | 0,973 |
| REM Latency |  |  |  |  |  |  | 0,003 | 0,002 | 0,135 | 0,002 | 0,002 | 0,381 |
| N3 % |  |  |  |  |  |  | -0,016 | 0,048 | 0,741 | 0,003 | 0,053 | 0,952 |
| N3 Latency |  |  |  |  |  |  | -0,003 | 0,004 | 0,339 | 0,003 | 0,005 | 0,596 |
| N2 % |  |  |  |  |  |  | -0,021 | 0,048 | 0,669 | 0,010 | 0,054 | 0,849 |
| N2 Latency |  |  |  |  |  |  | 0,008 | 0,013 | 0,536 | 0,018 | 0,016 | 0,271 |
| Awakenings |  |  |  |  |  |  | -0,002 | 0,014 | 0,898 | -0,001 | 0,016 | 0,958 |
| Delta Power |  |  |  |  |  |  |  |  |  | 0,367 | 0,707 | 0,604 |
| Sigma Power |  |  |  |  |  |  |  |  |  | 0,818 | 0,934 | 0,381 |
|  | Number of participants | | 246 | Number of participants | | 246 | Number of participants | | 246 | Number of participants | | 235 |
|  | Number of nights | | 1376 | Number of nights | | 1376 | Number of nights | | 1365 | Number of nights | | 1139 |
|  | Incremental R-squared | | 0,01 | Incremental R-squared | | 0,02 | Incremental R-squared | | 0,01 | Incremental R-squared | | 0,04 |

**Supplementary Table S2.** Within-participant effects on the subjective level of restedness. The table contains fixed effects associated with the deviation of objective sleep quality metrics from individual means. The table contains unstandardized regression coefficients, showing the expected change in restedness (in Likert points) as a function of a one-unit increase in sleep metrics. Sleep metrics are expressed as percentage points for sleep efficiency and sleep composition, minutes for total sleep time and sleep latency, total number for awakenings and log10 microvolt/sec^2^ for relative power. Incremental R^2^ refers to the variance accounted for by the models in addition to the variance accounted for by the random intercept and control variables. R^2^ values are shown for the full model, not only within-individual effects.

|  | Model 1 | | Model 2 | | Model 3 | | Model 4 | |
| --- | --- | --- | --- | --- | --- | --- | --- | --- |
|  | OR | p | OR | p | OR | p | OR | p |
| Sleep Efficiency | 1,080 | <0.001 |  |  |  |  |  |  |
| TST |  |  | 0,977 | <0.001 | 0,980 | 0,030 | 0,980 | 0,084 |
| SOL |  |  | 1,001 | 0,458 | 1,001 | 0,794 | 0,999 | 0,717 |
| WASO |  |  | 0,994 | 0,503 | 0,997 | 0,735 | 0,998 | 0,886 |
| REM % |  |  |  |  | 1,110 | 0,222 | 1,165 | 0,135 |
| REM Latency |  |  |  |  | 1,002 | 0,572 | 1,006 | 0,248 |
| N3 % |  |  |  |  | 1,099 | 0,266 | 1,134 | 0,209 |
| N3 Latency |  |  |  |  | 0,996 | 0,762 | 0,998 | 0,896 |
| N2 % |  |  |  |  | 1,080 | 0,380 | 1,128 | 0,244 |
| N2 Latency |  |  |  |  | 1,035 | 0,398 | 1,024 | 0,604 |
| Awakenings |  |  |  |  | 1,015 | 0,539 | 1,028 | 0,330 |
| Delta Power |  |  |  |  |  |  | 3,024 | 0,211 |
| Sigma Power |  |  |  |  |  |  | 0,208 | 0,084 |

**Supplementary Table S3.** Between-participant effects on subjectively rated deep sleep. The table contains fixed effects associated with the individual means of subjective sleep quality. The table presents odds ratios, indicating the impact of a one-unit change of sleep metrics on the odds of a response indicating poor sleep based on first GSQS item. Sleep metrics are expressed as percentage points for sleep efficiency and sleep composition, minutes for total sleep time and sleep latency, total number for awakenings and log10 microvolt/sec^2^ for relative power.

|  |  | Model 1 |  |  | Model 2 |  |  | Model 3 |  |  | Model 4 |  |
| --- | --- | --- | --- | --- | --- | --- | --- | --- | --- | --- | --- | --- |
|  | B | SE | p | B | SE | p | B | SE | p | B | SE | p |
| Sleep Efficiency | -0,034 | 0,020 | 0,085 |  |  |  |  |  |  |  |  |  |
| TST |  |  |  | 0,008 | 0,007 | 0,243 | 0,012 | 0,009 | 0,189 | 0,012 | 0,011 | 0,275 |
| SOL |  |  |  | 0,000 | 0,002 | 0,860 | 0,001 | 0,002 | 0,713 | 0,000 | 0,002 | 0,928 |
| WASO |  |  |  | 0,003 | 0,010 | 0,797 | -0,001 | 0,010 | 0,946 | -0,003 | 0,011 | 0,802 |
| REM % |  |  |  |  |  |  | 0,015 | 0,085 | 0,858 | 0,015 | 0,095 | 0,876 |
| REM Latency |  |  |  |  |  |  | -0,005 | 0,004 | 0,247 | -0,005 | 0,005 | 0,312 |
| N3 % |  |  |  |  |  |  | 0,034 | 0,084 | 0,686 | 0,045 | 0,093 | 0,628 |
| N3 Latency |  |  |  |  |  |  | 0,006 | 0,012 | 0,636 | 0,010 | 0,014 | 0,470 |
| N2 % |  |  |  |  |  |  | 0,029 | 0,086 | 0,734 | 0,044 | 0,096 | 0,647 |
| N2 Latency |  |  |  |  |  |  | 0,027 | 0,039 | 0,498 | 0,025 | 0,042 | 0,559 |
| Awakenings |  |  |  |  |  |  | -0,004 | 0,025 | 0,879 | 0,003 | 0,027 | 0,918 |
| Delta Power |  |  |  |  |  |  |  |  |  | 0,661 | 0,848 | 0,436 |
| Sigma Power |  |  |  |  |  |  |  |  |  | -1,194 | 0,874 | 0,172 |

**Supplementary Table S4.** Between-participant effects on the subjective level of restedness. The table contains fixed effects associated with the individual means of objective sleep quality, regressed on subjective rated restedness. The table contains unstandardized regression coefficients, showing the expected change in the level of restedness (in Likert points) as a function of a one-unit increase in sleep metrics. Sleep metrics are expressed as percentage points for sleep efficiency and sleep composition, minutes for total sleep time and sleep latency, total number for awakenings and log10 microvolt/sec^2^ for relative power.
